# Nociceptor-restricted cannabinoid receptor 1 contributes to chronic but not acute analgesia

**DOI:** 10.64898/2026.08.12.744456

**Authors:** Amber L. Milligan, Audrey R. Green, Katherine M. Garner, Thomas A. Szabo-Pardi, Luz R. Barron, Dana M. Jenkins, Carlos M. Castorena, Joel K. Elmquist, Michael D. Burton

## Abstract

Understanding the complex network that regulates pain is fundamental to develop strategies to combat its growing prevalence and increase useful therapeutics. Although extensive literature identifies the importance of cannabinoid receptors and endocannabinoids in controlling pain, their efficacy and loci of action remain debated. To directly test the actions of peripherally restricted cannabinoids and elucidate the minimal circuitry capable of producing cannabinoid-mediated analgesia, we utilized a novel genetic approach that allows for cell-specific reactivation of cannabinoid receptor 1 (CB1R) selectively in peripheral sensory neurons using newly developed CB1R floxed-stop-floxed mice (CB1R^LOXTB^) crossed with Nav1.8-cre mice (Na_v_1.8^+/-^:CB1R^LOXTB^). *Ex vivo* and *in vivo* experiments confirmed successful knockout and reactivation of CB1R. Wildtype littermate controls, but neither Na_v_1.8^+/-^:CB1R^LOXTB^ nor CB1R^LOXTB^ animals, exhibited robust analgesia after systemic WIN55,212-2 (WIN) treatment in the tail flick assay. Furthermore, the presence of CB1R on Na_v_1.8 neurons was not associated with either a difference in the development of inflammatory pain or the response to WIN. However, after neuropathic injury, CB1R^LOXTB^ animals displayed an earlier onset of both mechanical and thermal hypersensitivity than their Na_v_1.8^+/-^:CB1R^LOXTB^ or wildtype counterparts, suggesting a dual role for CB1R in inflammatory and neuropathic pain. These studies represent an important approach to further improve our mechanistic understanding of cannabinoid modulation of pain in the nervous system and begins to settle long-standing controversies in cannabinoid literature.

**Table of Contents:** Peripherally restricted cannabinoids show strong preclinical analgesic efficacy but have not translated clinically. Using a genetic model restricting CB1R to Nav1.8-expressing sensory neurons, we show peripheral neuronal endocannabinoid signaling is required for chronic, but not acute pain modulation. This dissociation suggests clinical failures may reflect testing peripheral cannabinoids in acute rather than chronic pain paradigms, informing future translational strategies.

## 1 Introduction

Pain is a complex sensory experience that affects more than 20% of adults worldwide every year, creating not only a major socioeconomic problem, but also a major quality of life disruption [1]. While in the United States, opioids are the predominant medication used to treat pain states, 20-30% of users become addicted, and accidental overdoses have risen exponentially since the late 1990s [2]. As both chronic pain and the opioid epidemic continue to plague the United States, it is imperative to discover effective alternative treatments.

Cannabinoids are thought to have low abuse potential and have been shown to have high analgesic efficacy in animal models, making them a potential novel therapeutic; however, these neuromodulators are highly debated [3, 4], in part because cannabinoid receptor 1 (CB1R) is one of the most widely expressed throughout the nervous system [5, 6], affecting many neurological and cognitive processes [7]. While global CB1R knockout studies have demonstrated the necessity of CB1R activation for both endogenous and exogenous pain inhibition [8, 9], these studies have not distinguished the specific neuroanatomical loci of action necessary for analgesia [10–15]. Moreover, it is unclear how CB1R modulates analgesia in the peripheral nervous system (PNS) versus in the central nervous system (CNS) [3, 16]. Furthermore, there is opposing evidence regarding the sufficiency of peripheral cannabinoid modulation, with some studies suggesting the requirement of CB1R in the CNS for analgesic effects [3, 17–20]. Agonism of CB1R in the CNS leads to several undesirable effects, including psychoactive reactions, impaired memory, anxiety, and impaired motor function [21–24]. This creates the need for peripherally restricted CB1R agonists to alleviate pain states [25–29]. However, in order to properly study the effects of CB1R on nociceptive processes, it is imperative that the effects in the PNS be distinct from the effects in the CNS.

The present study aims to investigate the specific role of CB1R on peripheral sensory neurons in anti-nociception by utilizing a novel mouse model that restricts CB1R expression to peripheral sensory neurons. In this model, a transcriptional blocker (TB) is flanked by lox-p sites, which can be removed in a cre-lox dependent manner. Thus, in the absence of cre, CB1R expression is blocked in the whole organism (CB1R^LOXTB^) [30]; when cre is expressed, CB1R is selectively expressed by only cre-containing cells. By crossing animals homozygous for the transcriptional blocker with animals expressing Na_v_1.8cre, a marker of peripheral sensory neurons, we are thus able to selectively re-express CB1R in peripheral sensory neurons, and separate the effects of cannabinoids in the PNS from the effects of cannabinoids in the CNS. In order to test the functionality of this model, we utilized several methods, both *in vivo* and *ex vivo*: CB1R-mediated cAMP decrease [31], gut motility via CB1R actions in gut innervating sensory neurons [32–34], and TRPV1 activity, as CB1R has been shown to modulate TRPV1-activity in peripheral sensory neurons [35].

Using our selective re-expression model, we hypothesized that CB1R action specifically on peripheral sensory neurons would be sufficient in producing analgesia. We expected that mice with CB1R re-expression only in peripheral sensory neurons (Na_v_1.8^+/-^:CB1R^LOXTB^) would have similar anti-nociceptive activity as their wild type (WT) counterparts. As cannabinoids have been shown to contribute to several different pain models, we elected to test our hypothesis in both inflammatory pain [36], including neurogenic inflammation, and neuropathic pain [37]. In addition to testing endogenous CB1R modulation of nociception, we also used cannabinoid agonist WIN55,212-2 (WIN) [38] in combination with our CB1R^LOXTB^ animals to test the effects of exogenous CB1R cannabinoid agonism.

## 2 Results

### 2.1 Generation of CB1R^LOXTB^ and Na_v_1.8^+/-^:CB1R^LOXTB^ animals

To validate our model, we utilized RNAscope *in situ* hybridization to show cell-specific knockdown and reactivation of *Cnr1* mRNA in the lumbar (L4-6) DRGs from WT, CB1R^LOXTB^, Na_v_1.8^+/-^:CB1R^LOXTB^, and ZP3^+/-^:CB1R^LOXTB^ animals (Fig. 1A). *Cnr1* mRNA was present and colocalized with *Scn10a* in WT animals and was significantly reduced in CB1R^LOXTB^ animals (Fig. 1B). For Na_v_1.8^+/-^:CB1R^LOXTB^ animals, *Cnr1* mRNA colocalized with cells containing *Scn10a* mRNA (Fig. 1A). These data provide further confirmation for the specificity and success of the model used going forward.

**Fig. 1:**
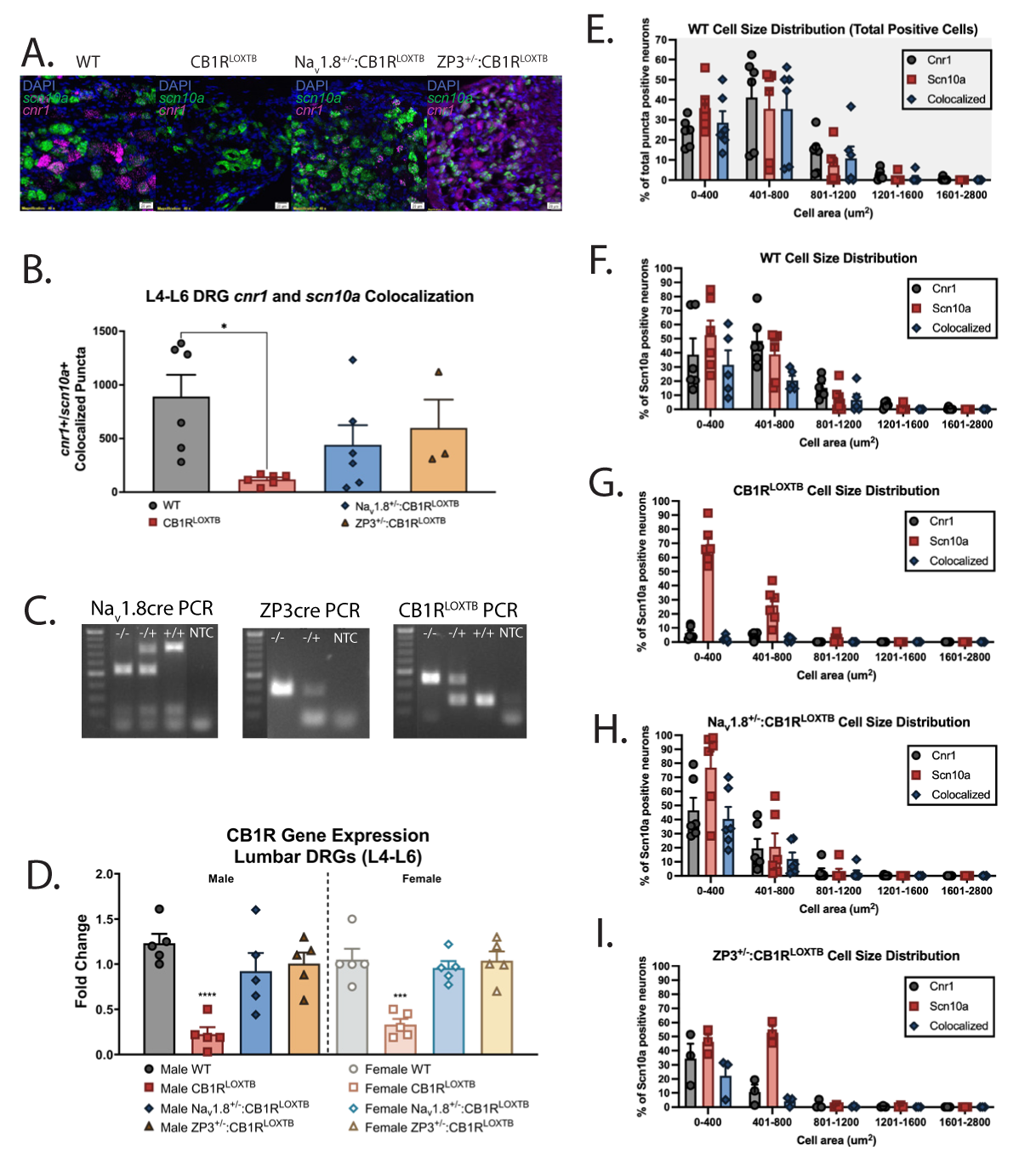
Generation of CB1R^LOXTB^, Na_v_1.8^+/-^:CB1R^LOXTB^, ZP3^+/-^:CB1R^LOXTB^ animals. (A) RNAscope i*n situ* hybridization of *Cnr1* (magenta) and *Scn10a* (green) mRNA in the lumbar (L4-6) DRGs. Scale bar = 20 μm. (B) Average colocalized *Cnr1*and *Scn10a* puncta (Tukey’s multiple comparisons test; n=3-6 animals, \**p*<0.05 WT vs. CB1R^LOXTB^). (C) PCR for Na_v_1.8Cre, ZP3Cre, and CB1R-TB. For each gene, a – indicates a WT mice, a ^-/+^ indicates heterozygous Na_v_1.8cre/ZP3cre/CB1R-TB, a ^+/+^ indicates homozygous Na_v_1.8cre/CB1R-TB. (D) Gene expression revealed insertion of a TB resulted in significant reduction of CB1R gene expression in both male and female lumbar DRGs. Crossing with either Na_v_1.8cre (cell specific) or ZP3cre (whole-body) restored CB1R expression to WT levels (Sidak’s multiple comparisons test; n=5 animals, 3 DRGs each, \*\*\*\**p*<0.0001 WT vs. CB1R^LOXTB^). (E) Cell size distribution of *Cnr1*, *Scn10a*, and colocalized puncta positive cells in WT lumbar DRGs. Cell size distribution of *Cnr1*, *Scn10a*, and colocalized *Scn10a* puncta positive cells in WT (F), CB1R^LOXTB^ (G), Na_v_1.8^+/-^:CB1R^LOXTB^ (H), and ZP3^+/-^:CB1R^LOXTB^ (I) lumbar DRGs.

Genotypes of Na_v_1.8^+/-^:CB1R^LOXTB^ and ZP3^+/-^:CB1R^LOXTB^ mice were confirmed with polymerase chain reaction (PCR) and gel electrophoresis (Fig. 1C). Real-time quantitative PCR was performed to validate successful knockdown and reactivation of CB1R transcript in Na_v_1.8-containing neurons via the transcriptional blocker (TB) model (Fig. 1D). CB1R^LOXTB^ animals expressed significantly less *Cnr1* mRNA in the DRG than WT animals (p<0.0001 for males; p=0.0007 for females; Table 2). WT, Nav1.8^+/-^:CB1R^LOXTB^, and ZP3^+/-^:CB1R^LOXTB^ DRGs expressed similar amounts of *Cnr1*. No sex difference in mRNA expression was observed; hence, validation data have been pooled between males and females.

Analysis of cell size distributions in WT lumbar DRGs depicted *Cnr1* puncta across a range of cell sizes and was colocalized with *Scn10a* in small and medium diameter cells (Fig. 1E). When comparing the cell sizes of *Scn10a* across WT, CB1R^LOXTB^, Na_v_1.8^+/-^:CB1R^LOXTB^, and ZP3^+/-^:CB1R^LOXTB^ DRGs, no differences were observed, so we can conclude the genetic modification was specific to CB1R (data not shown). Thus, to understand changes in size distribution relevant to the Na_v_1.8cre model, we normalized the cell counts out of *Scn10a* positive neurons. Cell size distributions in Na_v_1.8^+/-^:CB1R^LOXTB^ and ZP3^+/-^:CB1R^LOXTB^ DRGs were similar to the WT group, though Na_v_1.8^+/-^:CB1R^LOXTB^ *Cnr1* puncta was found in small and medium diameter neurons (Fig 1. F, H, I). *Cnr1* and colocalized *Cnr1* and *Scn10a* puncta positive cells were reduced in CB1R^LOXTB^ DRGs (Fig. 1G).

To assess CB1R functionality, we utilized the phenomenon where CB1R activation decreases forskolin-induced cAMP accumulation *in vitro* (Fig. 2A)[39]. As predicted, WIN treatment reduced cAMP in dissociated DRG neurons from WT animals (p<0.0001) but not CB1R^LOXTB^ animals; Na_v_1.8^+/-^:CB1R^LOXTB^ neurons recapitulated the WT phenotype (p<0.0001; Table 3). An *in vivo* validation experiment utilized the sensory neuron-mediated effects of CB1R to reduce gut motility [32]. As a subpopulation of Na_v_1.8-containing neurons are present in the nodose ganglia of the vagus nerve [40], this allows us to confirm functionality of our model independent of nociception. CB1R stimulation reduced gut motility in WT and Na_v_1.8^+/-^:CB1R^LOXTB^ animals, but not CB1R^LOXTB^ animals (p<0.0001; Fig. 2B; Table 4).

**Fig. 2:**
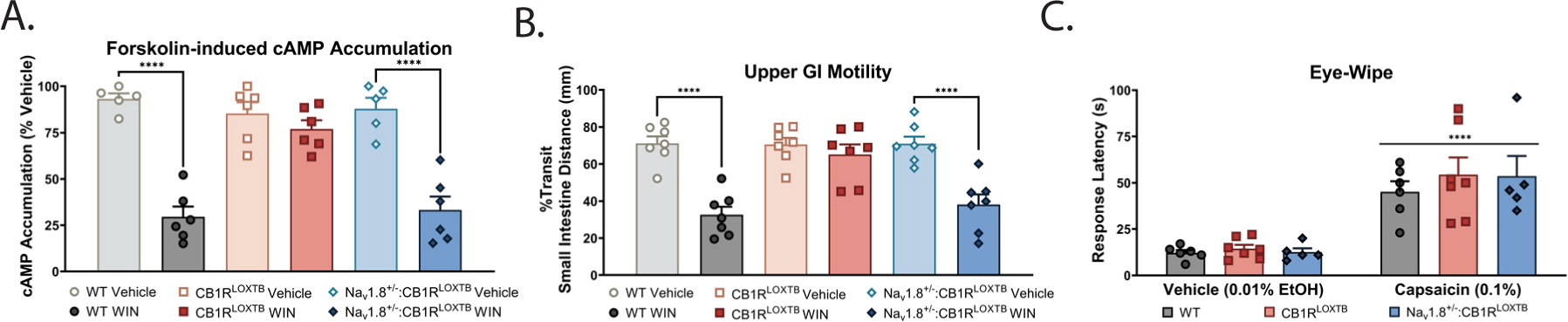
Functional Validation of CB1R^LOXTB^ and Na_v_1.8^+/-^:CB1R^LOXTB^ animals. (A) Reactivation of CB1R in Na_v_1.8 containing neurons restored a WT response to forskolin (Sidak’s multiple comparisons test; n=5-6 animals, \*\*\*\**p*<0.0001 WT VEH vs. WT WIN and Na_v_1.8^+/-^:CB1R^LOXTB^ VEH vs. Na_v_1.8^+/-^:CB1R^LOXTB^ WIN). (B) CB1R agonism reduced sensory neuron-mediated gut motility in WT and Na_v_1.8^+/-^:CB1R^LOXTB^ animals, but not CB1R^LOXTB^ animals. (Sidak’s multiple comparisons test; n=7 animals, \*\*\*\**p*<0.0001 WT VEH vs. WT WIN and Na_v_1.8^+/-^:CB1R^LOXTB^ VEH vs. Na_v_1.8^+/-^:CB1R^LOXTB^ WIN). (C) TRPV1 maintained functionality in CB1R^LOXTB^, Na_v_1.8^+/-^:CB1R^LOXTB^, and WT animals (ordinary two-way ANOVA; n=5-7 animals, \*\*\*\**p*<0.0001 WT VEH vs. WT capsaicin, CB1R^LOXTB^ VEH vs. CB1R^LOXTB^ capsaicin, Nav1.8^+/-^:CB1R^LOXTB^ VEH vs. Na_v_1.8^+/-^:CB1R^LOXTB^ capsaicin).

To understand if endogenous signaling was altered in our genetic model, we tested *in vivo* TRPV1 activity. TRPV1 and CB1R are known to highly colocalize in Na_v_1.8-containing neurons in mice [41], and CB1R knockout animals have been shown to have a blunted response to TRPV1 agonist capsaicin [42]. We therefore tested the animals’ responsivity to capsaicin in the eyewipe test (Fig. 2C). Capsaicin increased eyewipe time in all genotypes (p<0.0001; Table 5), suggesting our deletion was specific to CB1R and avoided off-target effects of CB1R knockout.

### 2.2 CB1R restricted to peripheral nociceptors does not mediate inflammatory pain nor reflexive analgesia

Models of inflammatory pain have been shown to engage various neuroimmune circuits [43]; as such, we elected to use intraplantar injection of carrageenan to investigate how CB1R in peripheral nociceptors mediates either a painful or inflammatory response. To assess endogenous cannabinoid modulation of inflammatory pain, 0.5% carrageenan was injected into the hindpaw to induce an inflammatory reaction and mechanical hypersensitivity and thermal hyperalgesia were recorded (Fig. 3A). WT, CB1R^LOXTB^, and Na_v_1.8^+/-^:CB1R^LOXTB^ animals developed similar mechanical hypersensitivity and thermal hyperalgesia in response to intraplantar carrageenan (Fig. 3B-C).

**Fig. 3:**
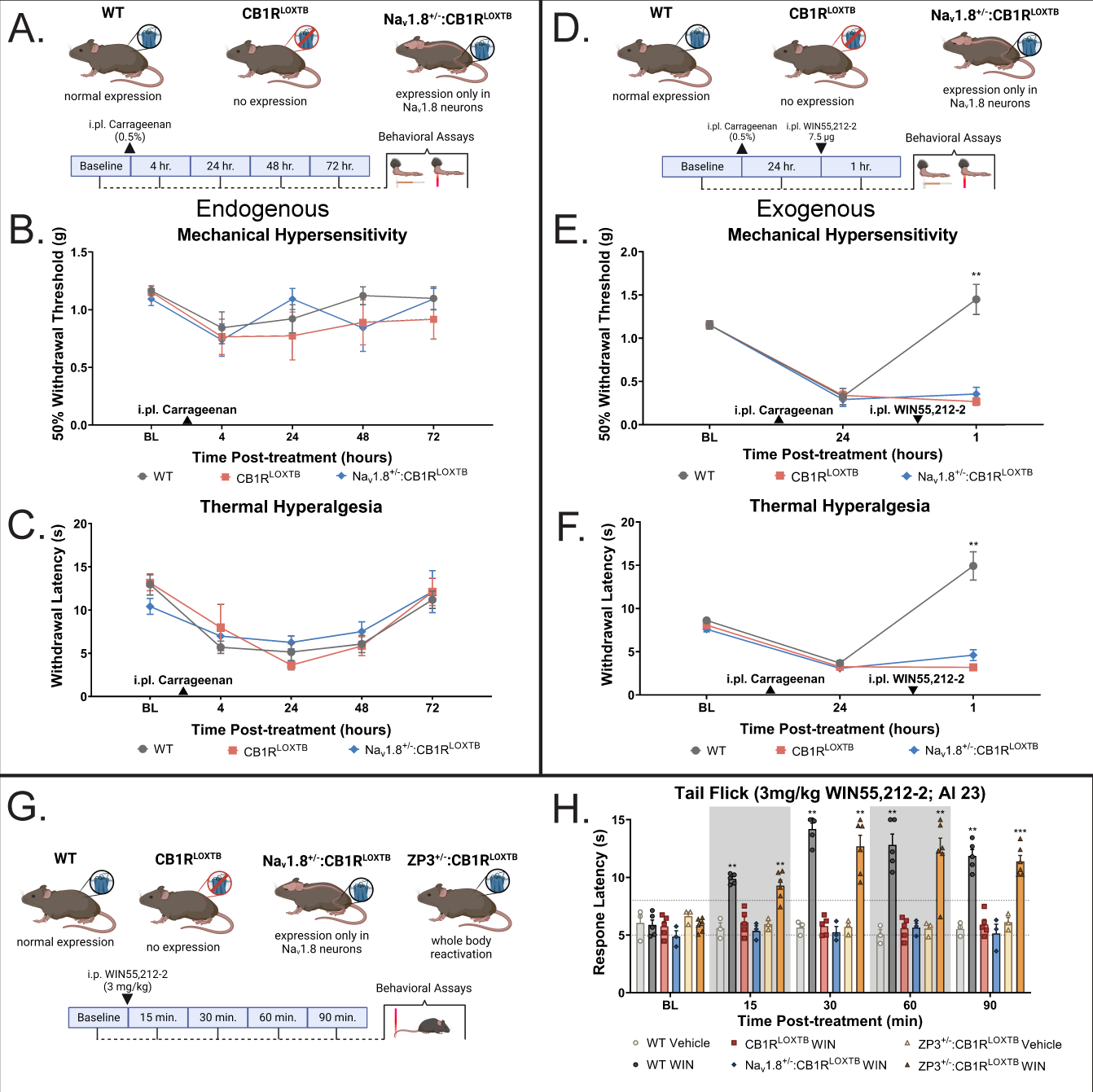
CB1R restricted to peripheral nociceptors does not mediate inflammatory pain nor reflexive analgesia. (A) Experimental schematic. WT, CB1R^LOXTB^, and Na_v_1.8^+/-^:CB1R^LOXTB^animals all had local inflammation induced with 0.5% intraplantar carrageenan; contralateral paw was used as control. Mechanical hypersensitivity and thermal hyperalgesia were recorded at the timepoints shown. (B) WT, CB1R^LOXTB^, and Na_v_1.8^+/-^:CB1R^LOXTB^ animals developed similar mechanical hypersensitivity in response to intraplantar carrageenan (Tukey’s multiple comparison test; n=5-7 animals). (C) WT, CB1R^LOXTB^, and Na_v_1.8^+/-^:CB1R^LOXTB^ animals developed similar thermal hyperalgesia in response to intraplantar carrageenan (Tukey’s multiple comparison test; n=5-7 animals). (D) Experimental schematic. WT, CB1R^LOXTB^, and Na_v_1.8^+/-^:CB1R^LOXTB^ animals all had local inflammation induced with 0.5% intraplantar carrageenan and were given 7.5 μg WIN or vehicle injected intraplantarly; contralateral paw was used as control. Mechanical hypersensitivity and thermal hyperalgesia were recorded at the timepoints shown. (E) WIN reduced carrageenan-induced mechanical hypersensitivity in WT but not CB1R^LOXTB^ or Na_v_1.8^+/-^:CB1R^LOXTB^ (Tukey’s multiple comparison test; n=5-7 animals, \*\**p*<0.01, WT vs. CB1R^LOXTB^ or Na_v_1.8^+/-^:CB1R^LOXTB^). (F) Local WIN caused analgesia in response to noxious heat in WT animals only (Tukey’s multiple comparison test; n=5-7 animals, \*\**p*<0.01, WT vs. CB1R^LOXTB^ or Na_v_1.8^+/-^:CB1R^LOXTB^). (G) Experimental schematic. WT, CB1R^LOXTB^, Na_v_1.8^+/-^:CB1R^LOXTB^, and ZP3^+/-^:CB1R^LOXTB^ animals were all given 3mg/kg WIN or its vehicle intraperitoneally and reflexive analgesia was recorded at the time points shown. (H) A high dose of WIN (3mg/kg) induced analgesia in WT and ZP3^+/-^:CB1R^LOXTB^ (whole-body reactivated) animals but did not induce analgesia in either CB1R^LOXTB^ or Na_v_1.8^+/-^:CB1R^LOXTB^ animals (Dunnett’s multiple comparisons test; n=3-6 animals, \*\*\**p*<0.001, \*\**p*<0.01, \**p*<0.05 vs. baseline). Dotted lines on graph (H) indicate baseline values (5-8 seconds). Grey boxes indicate 15- and 60-minute time points (H). Upward arrows indicate carrageenan administration (B, C, E, F); downward arrows indicate WIN administration (E and F).

CB1R agonists have been found to reduce inflammatory pain, both peripherally and centrally [44]. After inducing an inflammatory reaction as previously stated, 7.5 μg WIN was injected intraplantarly and mechanical hypersensitivity and thermal hyperalgesia were recorded (Fig. 3D). WIN reduced the carrageenan-induced mechanical hypersensitivity in WT animals but not CB1R^LOXTB^ and Na_v_1.8^+/-^:CB1R^LOXTB^ (Fig. 3E). Furthermore, WIN caused analgesia in response to noxious heat in WT animals only (p<0.01; Fig. 3F).

As literature does not suggest sex differences in CB1R activity and we found no sex differences in WIN-mediated analgesia at either a low (1mg/kg) or high (3mg/kg) dose (Fig. S1A; p<0.0001, Tables 6-7), male and female data have henceforth been pooled. We failed to recapitulate a WT phenotype in Na_v_1.8^+/-^:CB1R^LOXTB^ at either a low (Fig. S1B; Table 8) or high (Fig. S1C; Table 9) dose of WIN. Whole-body reactivation in ZP3^+/-^:CB1R^LOXTB^ restored the analgesic properties of WIN (Fig. 3G and H; Table 10); however, CB1R^LOXTB^ and Na_v_1.8^+/-^:CB1R^LOXTB^ animals consistently did not respond to WIN (Fig. 3G and H; Table 10). However, as tail flick behavior is mediated by a spinal circuit and CB1R is also expressed on interneurons in the spinal cord [45], WIN may exert analgesic effects by acting on these cells. Most importantly, these data suggest that reflexive pain is modulated by the spinal cord rather than peripheral circuits. Thus, to solidify the role of peripheral CB1R in analgesia, our later experiments tested peripheral insults.

### 2.3 CB1R deleted from peripheral nociceptors does not mediate inflammatory pain nor reflexive analgesia, but may modulate neurogenic inflammation

Previous studies have utilized a flox model, where CB1R is only removed from Na_v_1.8 containing neurons, to suggest CB1R in peripheral nociceptors is necessary to mediate an analgesic response [41]. As our previous experiments suggested CB1R in peripheral nociceptors is not required for analgesia, we initially explored if animals with CB1R either present in whole-body (CB1R^fl/fl^) or removed from only Na_v_1.8 containing peripheral nociceptors (Na_v_1.8^+/-^:CB1R^fl/fl^) responded differently to inflammatory pain (Fig. S2A-D). Local carrageenan induced inflammatory pain in both genotypes (Fig. 4A-C; Tables 11-14). Interestingly, systemic WIN (1mg/kg) both reversed mechanical hypersensitivity (Fig. 4B and S3A; Tables 11-12) and reduced thermal hyperalgesia independent of the presence of CB1R on Na_v_1.8 expressing neurons (Fig. 4C and S3A; Tables 13-14). This suggests that other cell types are responsible for CB1R-mediated analgesia. However, from this data alone, it is not clear if the effects of CB1R signaling are due to peripheral cells (Schwann cells, myeloid cells) or central activity (microglia, spinal interneurons, CB1R in the cortex or periaqueductal grey).

**Fig. 4:**
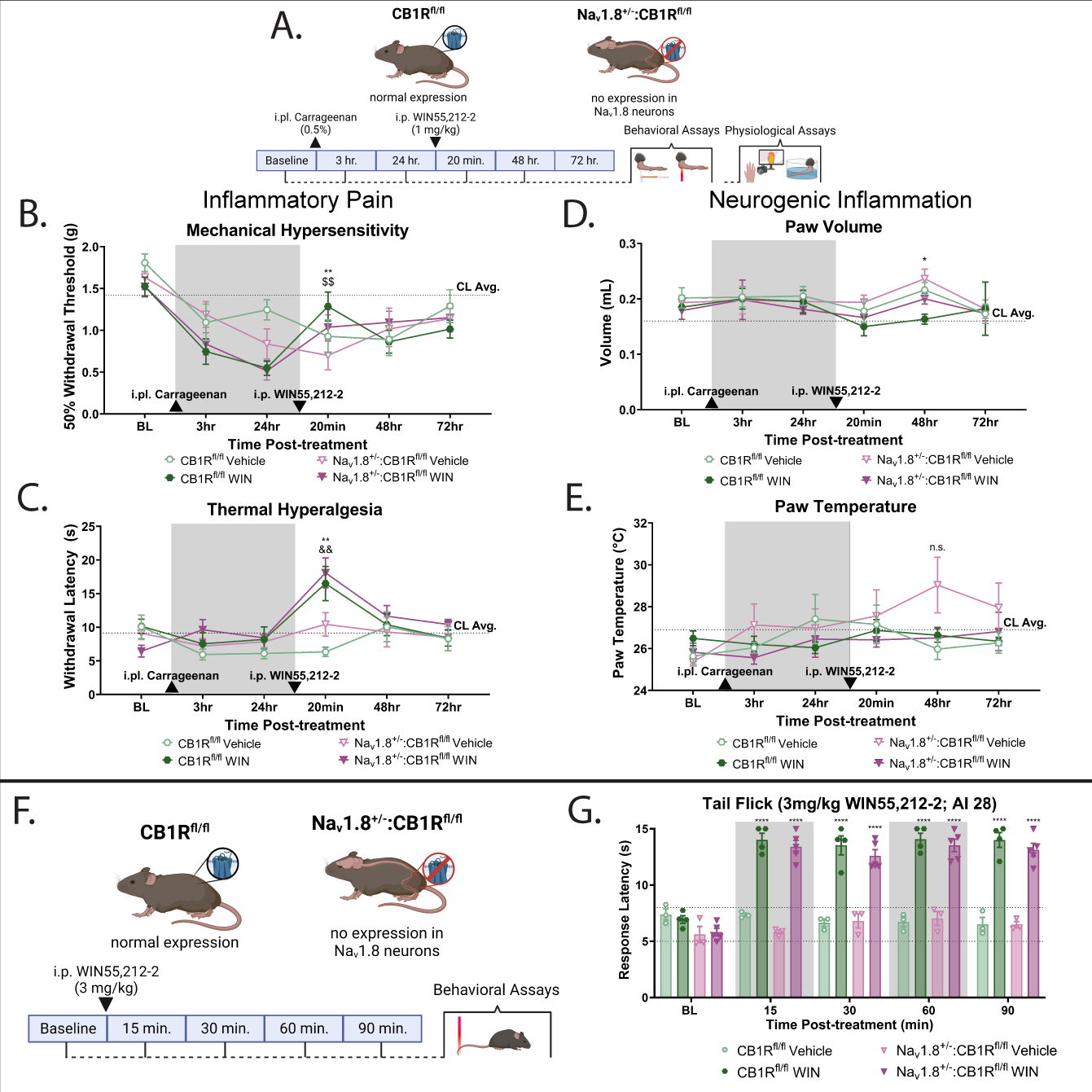
CB1R deleted from peripheral nociceptors does not mediate inflammatory pain nor reflexive analgesia, but may modulate neurogenic inflammation. (A) Experimental schematic. Note that rather than using TB animals, CB1R^fl/fl^ animals were used; thus, CB1R deletion is limited to Na_v_1.8-expressing tissues. Animals all had local inflammation induced with 0.5% intraplantar carrageenan and were given 1mg/kg WIN or its vehicle intraperitoneally; contralateral paw was used as control. Mechanical hypersensitivity, thermal hyperalgesia, paw volume, and paw temperature were recorded at the timepoints shown. (B) CB1R^fl/fl^ and Na_v_1.8^+/-^:CB1R^fl/fl^ animals both developed mechanical hypersensitivity after carrageenan. Both genotypes experienced analgesia in response to systemic WIN (Tukey’s multiple comparisons test, n=8-12 animals, \*\**p*<0.01 CB1R^fl/fl^ vehicle vs CB1R^fl/fl^ WIN, $$*p*<0.01 Na_v_1.8^+/-^:CB1R^fl/fl^ vehicle vs. Na_v_1.8^+/-^:CB1R^fl/fl^ WIN). (C) WIN reversed carrageenan-induced thermal hyperalgesia in both CB1R^fl/fl^ and Na_v_1.8^+/-^:CB1R^fl/fl^ animals (Tukey’s multiple comparisons test, n=8-12 animals, \*\**p*<0.01 CB1R^fl/fl^ vehicle vs CB1R^fl/fl^ WIN, &&*p*<0.01 Na_v_1.8^+/-^:CB1R^fl/fl^ vehicle vs. Na_v_1.8^+/-^:CB1R^fl/fl^ WIN). (D) WIN blocked carrageenan-mediated inflammation in CB1R^fl/fl^ animals only (Tukey’s multiple comparisons test, n=8-12 animals, \**p*<0.05 CB1R^fl/fl^ vehicle vs CB1R^fl/fl^ WIN). (E) No differences in paw temperature were seen across genotypes or treatment. (Tukey’s multiple comparisons test, n=8-12 animals). Dotted line indicates contralateral average (G) Experimental schematic. WT, CB1R^fl/fl^, and Na_v_1.8^+/-^:CB1R^fl/fl^ animals were all given 3mg/kg WIN or its vehicle intraperitoneally and reflexive analgesia was recorded at the time points shown. (H) A high dose of WIN (3mg/kg) induced analgesia in CB1R^fl/fl^ and Na_v_1.8^+/-^:CB1R^fl/fl^ animals (Dunnett’s multiple comparisons test; n=3-5 animals, \*\*\*\**p*<0.0001 vs. baseline). Dotted lines on graph (G) indicate baseline values (5-8 seconds). Grey boxes indicate 15- and 60-minute time points (G).Upward arrows indicate carrageenan administration and downward arrows indicate WIN administration (B, C, D, E).

Furthermore, carrageenan increased paw volume in both CB1R^fl/fl^ and Na_v_1.8^+/-^:CB1R^fl/fl^animals (Fig.4D; Table 15), but WIN blocked this increase only in CB1R^fl/fl^ animals. Similar trends were not seen in paw temperature data (Fig. 4E; Table 16). Based on data suggesting nociceptive neurons are capable of modulating inflammation after initial pain [46], our data suggest that CB1R activity in neurons may signal to other tissues to ultimately reduce inflammation.

After CB1R^fl/fl^ and Nav1.8^+/-^:CB1R^fl/fl^ animals received WIN in the tail flick assay (Fig. 4G and H), both genotypes experienced reflexive analgesia (Fig. 4H). This further supports the suggestion that other peripheral or central cell types are responsible for CB1R-mediated analgesia.

### 2.4 CB1R restricted to peripheral nociceptors mediates neuropathic pain

Although CB1R may not mediate analgesia or onset of pain after inflammatory insult, peripheral nerve injury has been shown to upregulate CB1R in the CNS [47, 48], although it is unclear how this correlates to peripheral nociceptors. Additionally, the role of peripheral nociceptors in CB1R’s ability to decrease in neuronal excitability remains to be elucidated. As such, we initiated one final test, hypothesizing that CB1R restricted to Na_v_1.8+ nociceptors is sufficient to cause pain relief after spared nerve injury (SNI) (Fig. 5A).

**Fig. 5:**
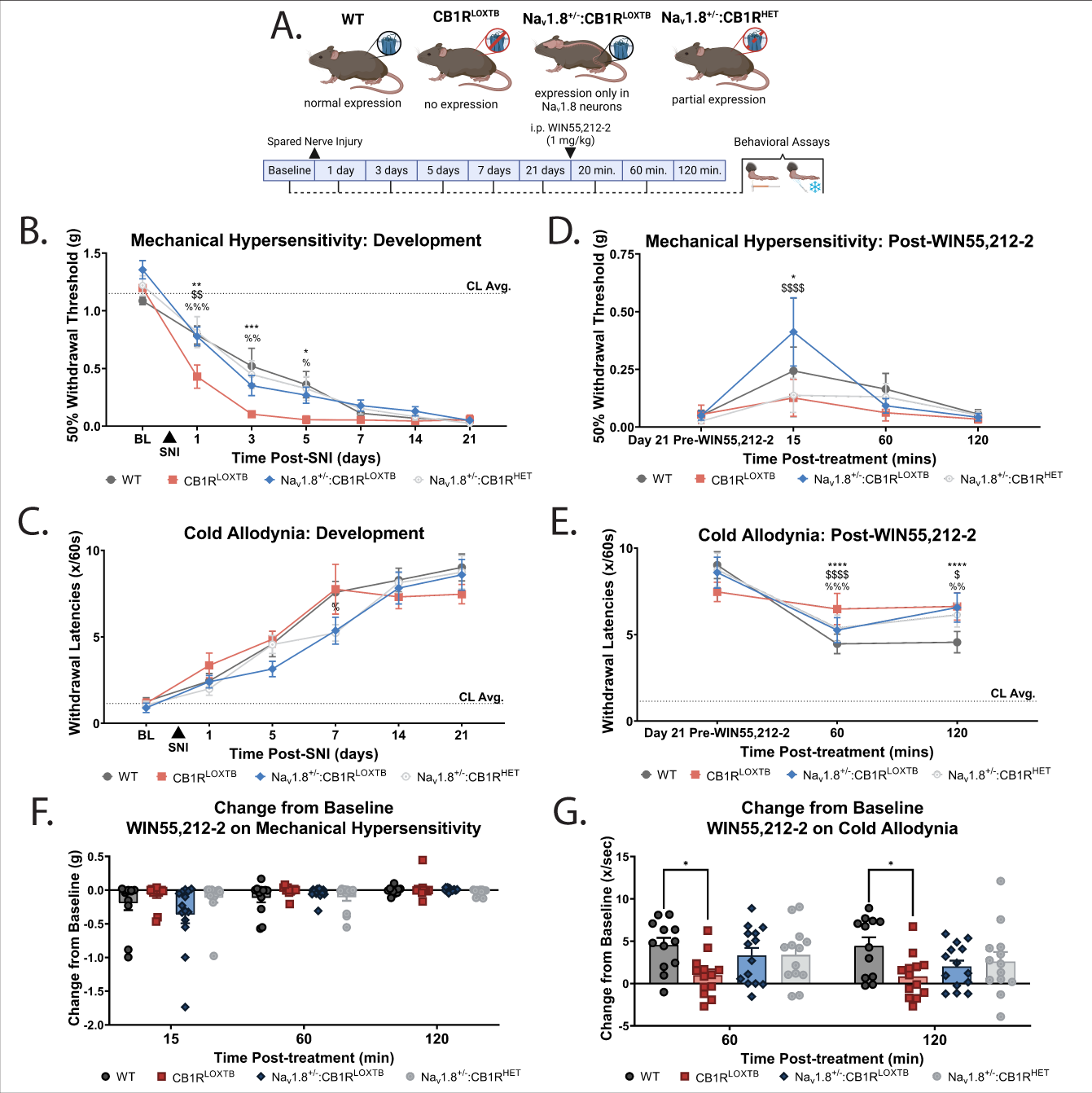
CB1R restricted to peripheral nociceptors mediates neuropathic pain. (A) Experimental Schematic. Animals expressing various levels of CB1R in peripheral DRGs were given SNI surgery to induce neuropathic pain; on day 21, 1mg/kg WIN was given and analgesia was tested at 20 minutes, 60 minutes, and 120 minutes. (B) WT, Na_v_1.8^+/-^:CB1R^LOXTB^, and Na_v_1.8^+/-^:CB1R^HET^ animals developed mechanical hypersensitivity later than CB1R^LOXTB^ animals (Bonferroni’s multiple comparisons test, n=12-13 animals, vs CB1R^LOXTB^: * WT, $ Na_v_1.8^+/-^:CB1R^LOXTB^, % Na_v_1.8^+/-^:CB1R^HET^, \*\*\**p*<0.001, \*\**p*<0.01, \**p*<0.05). (C) Animals developed cold allodynia similarly after SNI, although Na_v_1.8^+/-^:CB1R^HET^ animals were slightly less sensitive than CB1R^LOXTB^ counterpoints (Bonferroni’s multiple comparisons test, n=12-13 animals, %*p*<0.05 CB1R^LOXTB^ vs. Na_v_1.8^+/-^:CB1R^HET^). (D) WIN mediated analgesia to mechanical hypersensitivity in WT and Na_v_1.8^+/-^:CB1R^LOXTB^ only (Bonferroni’s multiple comparisons test, n=12-13 animals, vs BL: * WT, $ Nav1.8^+/-^:CB1R^LOXTB^, \*\*\*\**p*<0.0001, \**p*<0.05). (E) WIN only caused analgesia in response to cold stimuli when CB1R was present on peripheral nociceptors (Bonferroni’s multiple comparisons test, n=12-13 animals, vs BL: * WT, $ Na_v_1.8^+/-^:CB1R^LOXTB^, % Na_v_1.8^+/-^:CB1R^HET^, \*\*\*\**p*<0.0001, \*\*\**p*<0.001, \*\**p*<0.01, \**p*<0.05). Dotted line indicates sham average (n=17, mix of genotypes with no differences seen; all SNI animals significantly different from sham). (F) Change from baseline analysis of WIN on mechanical hypersensitivity (Tukey’s multiple comparison test, n=12-13 animals). (G) Change from baseline analysis of WIN on cold allodynia (Tukey’s multiple comparison test, n=12-13 animals).

CB1R^LOXTB^ animals experienced the onset of mechanical hypersensitivity more quickly after SNI than their wild type or Na_v_1.8^+/-^:CB1R^LOXTB^ counterparts (Fig. 5B; Table 21).

Interestingly, both wild type and Na_v_1.8^+/-^:CB1R^LOXTB^ animals sensitized similarly after SNI, with Na_v_1.8^+/-^:CB1R^HET^ animals behaving similarly to their wildtype and fully reactivated counterparts (Fig. 5B; Table 21). No differences in cold hypersensitivity were seen between groups (Fig. 5C; Table 22). All animals eventually developed similar hypersensitivities by day 7 (mechanical) or 14 (thermal) (Fig. 5B-C). These data suggest endocannabinoids modulate the onset, but not maintenance, of neuropathic pain.

Additionally, we assessed whether exogenous cannabinoids could induce analgesia after hypersensitivity had developed, or if cannabinoids were solely protective during the onset of neuropathy. Animals were treated with WIN 21 days after SNI, when all animals were at similar hypersensitivities (Fig. 5D-E; Table 23-24). Interestingly, WIN treatment was sufficient to increase the 50% mechanical withdrawal thresholds of both Na_v_1.8^+/-^:CB1R^LOXTB^ animals and wildtype animals but had no effect in CB1R^LOXTB^ or Na_v_1.8^+/-^:CB1R^HET^ animals (Fig. 5D, F; Table 23). However, only wildtype animals experienced a decrease in cold allodynia after WIN treatment (Fig. 5E, G; Table 24).

## 3 Discussion

In this study, we utilized a novel genetic model to restrict CB1R expression to peripheral sensory neurons. Contrary to existing literature [41], we did not find that CB1R restricted to Na_v_1.8-containing nociceptors was responsible for acute analgesia. Na_v_1.8^+/-^:CB1R^LOXTB^ that expressed CB1R only on peripheral nociceptors responded similarly to their whole-body knockout counterparts in both the tail flick assay (Fig. 3H) and in a model of inflammatory pain (Fig. 3A-F); Na_v_1.8^+/-^:CB1R^fl/fl^ animals that lacked CB1R in peripheral nociceptors experienced analgesia after WIN (Fig. 4B-C, H). However, Na_v_1.8^+/-^:CB1R^fl/fl^ animals did not experience the same decrease in local inflammation as their WT counterparts did (Fig. 4E), and Na_v_1.8^+/-^:CB1R^LOXTB^ animals behaved similarly to their WT counterparts after nerve injury (Fig. 5).

### 3.1 Generation of a novel model to study peripherally restricted CB1R

Our first major finding was the development of a genetic model to test specific CB1R activity in Na_v_1.8 containing neurons. Other studies have utilized peripherally restricted CB1R agonists [49]; however, some evidence suggests these compounds are capable of crossing the blood-brain barrier and exerting central effects [50]. The development of a genetic model elucidates the effects of CB1R in peripheral nociceptors while avoiding this potential (Fig. 1). The addition of a translational blocker (TB) to the genome was sufficient to reduce CB1R expression in DRG. Removal of this block via cre recombinase (both Na_v_1.8 cre and ZP3 cre) was able to induce the re-expression of CB1R in peripheral sensory neurons. Functionality of the reactivation was tested both *in vivo* via a gut motility assay and *ex vivo* via agonist-induced cAMP production. CB1R agonism has been shown to inhibit forskolin-stimulated cAMP production [31]. After forskolin stimulation, cultured DRG neurons from Na_v_1.8^+/-^:CB1R^LOXTB^animals had a decrease in cAMP comparable to wildtype animals, whereas CB1R^LOXTB^ animals did not exhibit a response to stimulation. Furthermore, *in vivo*, Na_v_1.8^+/-^:CB1R^LOXTB^ but not CB1R^LOXTB^ had reduced gut motility in response to a CB1R agonist, consistent with the role of vagal CB1R in gut motility [32–34]. Furthermore, TRPV1 activity was not affected by deletion or reactivation of CB1R, suggesting the genetic manipulation did not have off target effects.

A final establishing piece of data was CB1R agonism having equivalent effects in animals of both sexes (Fig. S1A). As males and females are thought to utilize different mechanisms in nociception [51], this data suggests that CB1R agonism could be an effective way of treating pain in both sexes. This is clinically relevant because females report pain more often than males [52], thus representing a need for effective sex-independent therapeutics.

### 3.2 CB1R in peripheral nociceptors plays a dual role in mediating nociception

Carrageenan alone was able to induce inflammation, marked by an increase in paw size and decrease in both mechanical and thermal withdrawal thresholds, independent of CB1R expression (Fig. 4D and E). However, WIN caused analgesia when CB1R was removed from only Na_v_1.8 containing neurons (Fig. 4); when CB1R was only present on Na_v_1.8 neurons, the drug had no effect (Fig. 3). Furthermore, WIN only reduced inflammation when CB1R was present on Na_v_1.8 containing neurons (Fig. 4D). This suggests CB1R in nociceptors may communicate to control a later inflammatory response, with its activity in other cells being responsible for pain relief [36]. Others have found a disconnect between pain and inflammation, that suggests the two are related but independent processes [46, 53, 54]. Studies that have looked at cannabinoid-mediated analgesia have shown that peripherally-restricted cannabinoids are effective in relieving inflammatory pain, albeit without testing inflammation itself [50].

Interestingly, CB1R on peripheral sensory neurons appears to modulate pain types differently. As discussed, CB1R on peripheral sensory neurons may play a role in inflammation, but not pain (Fig. 3). However, after nerve injury, CB1R^LOXTB^ animals exhibited sensitivity sooner than their wildtype or Na_v_1.8^+/-^:CB1R^LOXTB^ counterparts (Fig. 5B). One of the hallmarks of SNI is ectopic activity in the affected DRGs [55–57]. As CB1R is capable of decreasing neuronal excitability via the G_i_/G_o_ pathway, this suggests that nerve injury can endogenously drive the nervous system to modulate its own ectopic activity. Furthermore, exogenous CB1R agonism was sufficient to reduce mechanical hypersensitivity after neuropathic injury, although partial reactivation of CB1R was not sufficient to cause this phenotype. However, development of cold allodynia was not modulated by CB1R, and the presence of CB1R on Na_v_1.8+ neurons was not sufficient to cause a response to WIN. Although inflammation commonly accompanies neuropathic pain [58], and may be responsible for its initial response, changes to the circuitry in the sciatic nerve, its innervating DRGs, and the spinal cord are responsible for the maintenance of chronic neuropathic pain [59, 60]. As SNI induces ectopic activity in nociceptors, this suggests that the actions of CB1R in neuropathic pain are related to its functions in reducing neuronal excitability [61, 62].

However, it should be noted that CB1R has been shown to be highly expressed in large diameter sensory neurons that are typically proprioceptive [14]. These fibers not only express Na_v_1.8 [63], but are also implicated in circuits mediating post-SNI induction of allodynia, [64, 65]. Although we hypothesize our results were due to CB1R restricted to small diameter nociceptors, we did not distinguish the role of these large-diameter nociceptors. Future studies may elect to use a marker for these neurons, such as Pirt cre [66], to elucidate the contributions of small-diameter vs. large-diameter neurons to analgesia.

It has been suggested that receptors that have both anti-inflammatory and anti-nociceptive properties can mask which property is ultimately responsible for pain relief [67]. The present study suggests that CB1R in the periphery does utilize anti-inflammatory and anti-nociceptive signaling to ultimately cause analgesia. Our study provides a potential explanation for known issues in creating efficacious peripherally restricted CB1R analgesics [68]; restriction of cannabinoid activity to nociceptors is not necessary for pain relief, leading to conclude that other cell types, both peripheral and central, could be responsible for the analgesic effects of CB1R. As peripherally restricted cannabinoids can cross into the central nervous system with sufficient dosing [50, 68], it is still unclear how to best utilize peripherally restricted CB1R agonists as analgesics. Ultimately, further elucidating the cell types that are involved in the analgesic effects of CB1R will aid in advancing our mechanistic understanding of cannabinoid modulation of pain.

## 4 Materials and Methods

### 4.1 Animals

All animal experiments were carried out in accordance with protocols approved by the Institutional Animal Care and Use Committee of the University of Texas at Dallas. Mice were housed in a temperature and humidity-controlled facility maintained on a 12-hour light/dark cycle (lights on 0600h, lights off 1800h). Mice had ad-libitum access to food and water.

All animals used in experiments were bred in house; animals were weaned at 28 days, at which time a tail biopsy was taken and used for genotyping via PCR. Transgenic animals were backcrossed at least eight generations to maintain a C57BL/6 background. Eight to twelve-week-old adult mice (male, 25-30g; female 20-25g) were used for all behavioral and biochemical assays. CB1R^LOXTB^ mice (gift from JK Elmquist) were generated via the insertion of a transcriptional blocker (TB) inserted in the CB1R gene between the second and third exon. Lox-p sites were inserted before and after the TB, allowing for removal and CB1R expression in a cre-lox dependent manner [30]. Animals were crossed with Na_v_1.8cre animals (originally from Professor John Wood at University College London, commercially available from Intrafrontier, EMMA ID 04582, now available at Jackson strain # 036564), a Cre driver specifically chosen for its high efficiency [69]. This generated Na_v_1.8^+/-^:CB1R^LOXTB^ animals that only expressed CB1R in peripheral sensory neurons. Although both CB1R^LOXTB^ and Na_v_1.8^+/-^:CB1R^LOXTB^ animals were homozygous for the transcriptional blocker, in spared nerve injury experiments, animals heterozygous for both Na_v_1.8cre and the CB1R transcriptional blocker (Na_v_1.8^+/-^:CB1R^HET^). CB1R^LOXTB^ animals were also crossed with zona pellucida 3 (ZP3) cre (Jackson strain #003651) [70] to generate ZP3^+/-^:CB1R^LOXTB^ animals, functionally reactivating CB1R in whole-body. In some experiments, Na_v_1.8cre animals were crossed with CB1R^fl/fl^ animals (gift from JK Elmquist) rather than CB1R^LOXTB^ animals to generate Na_v_1.8^+/-^:CB1R^fl/fl^ animals with CB1R floxed from peripheral nociceptors [71]. C57BL/6 animals with no genetic modifications were used as wildtype controls. A summary of genotypes is contained in Table 1.

### 4.2 Polymerase Chain Reaction (PCR)

Animals were weaned between 21-28 days old, at which time a tail clip biopsy (∼1mm) was collected. DNA was extracted from tail clips using 75ul extraction buffer (25mM NaOH, 0.2 mM EDTA in ddH2O) and incubated at 98°C for one hour. An equal amount of neutralization buffer (40 mM Tris-HCl, pH 5.5) was added following incubation. DNA samples were stored at −20°C until genotyping. PCR was performed using JumpStart REDTaq Ready Mix (Sigma, Cat#P0982-800RXN) and the primers listed below [72]. Genotyping for Na_v_1.8cre, Zp3cre, CB1RTB, and CB1R-Flox were each performed as a single reaction. The CB1RTB gene was amplified using the primer set CB1RTB_Cb1pDis+ 5’-CAC TCT GAC TTT TCT TGC CAC TC-3’, CB1RTB_M274 5’-CAG ATG TAG CTA AAA GGC CTA TCA CAA ACT-3’, and CB1RTB_Ex2R2 5’-TGG TTG TGT CTC CTG CTG GAA-3’. The Zp3cre gene was amplified using the primer set IMR1084-Cre-Fwd 5’-GCG GTC TGG CAG TAA AAA CTA TC-3’, IMR1085-Cre-Rev 5’-GTG AAA CAG CAT TGC TGT CAC TT-3’, IMR7338-Control-Fwd 5’-CTA GGC CAC AGA ATT GAA AGA TCT-3’, and IMR7339-Control-Rev 5’-GTA GGT GGA AAT TCT AGC ATC ATC C-3’. The CB1R-Flox gene was amplified using the primer set FLXCB1R-1 5’-ACC ACC TTC CTC ATG TTA AAC T-3’, FLXCB1R-2 5’-GAC CAG AGA CAG CTC CAG A-3’, and FLXCB1R-3 5’-TGA GGG CTA TAT TCT GTT TTT GC-3’. The Na_v_1.8cre gene was amplified using the custom primer pair newNav-WT-Fwd 5’ GAT GGA CTG CAG AGG ATG GA-3’ and newNAV-Cre-Rev 5’-CGT ATA TCC TGG CAG CGA TC-3’. The Na_v_1.8 wild-type gene was amplified using the custom primer pair newNAV-WT-Fwd 5’ GAT GGA CTG CAG AGG ATG GA-3’ and newNAV-WT-Rev 5’-GGT GTG TGC TGT AGA AAG-3’. Samples were run on a 1.5% agarose (VWR, Cat# MPN605-500G) and 1X Tris-Acetate-EDTA (TAE) gel and imaged using a ChemiDoc (BioRad) and ImageLab. DNA Ladder (100bp, Thermo Scientific GeneRuler 100bp) was used for size determination. The Na_v_1.8cre and Na_v_1.8 WT bands are at 800 bp and 500 bp, respectively. The CB1RTB floxed and CB1RTB WT bands are at 200bp and 350bp, respectively. The CB1R-Flox floxed and CB1R-Flox WT bands are at 233bp and 195bp, respectively. The Zp3cre and Zp3 internal control are at 100bp and 350bp, respectively.

### 4.3 RNAscope In Situ Hybridization

RNA *in situ* hybridization was performed on fresh frozen DRGs sectioned at 14 μm using the RNAscope *in situ* hybridization assay (Multiplex Fluorescent Reagent Kit Version 2) with the Advanced Cell Diagnostics (ACD) protocol [72]. The following adjustments were made: samples were incubated in fresh 10% neutral buffered formalin (NBF) at 4°C for one hour and a 30 second Protease IV digestion was used. Akoya Biosciences fluorophores were used at a 1:1500 dilution, including TSA Plus Fluorescein (Cat# NEL741001KT) for C1 and TSA Plus Cyanine 5 (Cat# NEL745001KT) for C3. The probes used were Mm-Scn10a (Cat#426011) and Mm-Cnr1-C3 (Cat# 420721-C3). The RNAscope 3-plex negative control probe (Cat#320871), which targets the bacterial gene *DapB*, and the positive control probe that targets a highly expressed transcript (Ubiquitin C) in all cell types (Mm-UBC, Cat#310771) were used.

Representative images for publication were acquired using the Olympus FV300RS Confocal Laser Scanning Microscope using the 40× objective.

### 4.4 Cell Size Measurements

Analysis of cell size distributions were conducted using the Olympus CellSens Software. Utilizing the RNA *in situ* hybridization confocal images, cells were outlined and their area was measured [73]. Only cells with visible nuclei were included in the analysis.

### 4.5 Quantitative Real-Time PCR

Animals (8–16 weeks old) were deeply anesthetized with isoflurane and euthanized by cervical dislocation. Lumbar dorsal root ganglia (L4–6) were extracted from both sides of the spinal column and frozen in liquid nitrogen. Total RNA was extracted using RNA Stat60 (Teltest). cDNA was synthesized using the high-capacity cDNA Kit (Applied Biosystems).

TaqMan Primers for Cnr1 (CB1R) (ID:Mm01212171_s1) and 18 s (ID: Hs99999901_s1) were purchased from Applied Biosystems. The mRNA contents were normalized to 18 s mRNA levels. All assays were performed using an ABI Prism 7900HT sequence detection system (Applied Biosystems). The relative amounts of all mRNAs were calculated using the ΔΔCT assay to express fold change from WT group [74]. Experimenters were blinded to both sex and genotype of animals during data collection.

### 4.6 Drugs

Λ-Carrageenan (Cayman Chemical, Cat#22049) was injected intraplantarly at 0.5%; the contralateral hindpaw was used as naive control. Animals that were not statistically different from contralateral hindpaw (i.e. did not successfully receive carrageenan) were excluded from experiment. WIN55,212-2 21 (Cayman Chemical, Cat#10009023) was given intraperitoneally at a dose of 1 or 3 mg/kg or intraplanterly at 7.5 μg; vehicle was 30% DMSO in 1X PBS. In neurogenic inflammation experiments, vehicle was 30% DMF; no differences were determined between WIN made in DMF and DMSO or its equivalent vehicle.

### 4.7 Culture and cAMP Assay

Whole DRGs from male and female WT, CB1R^LOXTB^, and Na_v_1.8^+/-^:CB1R^LOXTB^ animals were aseptically removed and kept in ice-cold HBSS (Gibco, Cat#14-170-112). DRGs were enzymatically digested digestion in collagenase A (1mg/mL HBSS; Sigma-Aldrich, Cat#10103586001) for 20 minutes, followed by collagenase D (1mg/mL 10% papain in HBSS; Sigma-Aldrich, Cat#1188866001 and Cat#10108014001) for 20 minutes. Digestion was halted with trypsin inhibitors (1mg/mL 10% BSA in TG media, Sigma-Aldrich, Cat#10109886001). DRGs were then manually triturated and strained in a 70uM mesh strainer to separate individual neurons. Neurons were plated in TG media (10% fetal bovine serum (HyClone, Cat#SH30088.03), 1% penicillin/streptomycin (Fisher Scientific, Cat#15070063) in DMEM (HyClone, Cat#SH30023)) on sterile poly-D lysine coated glass coverslips (Sigma-Aldrich, Cat#P0899) in a 5% CO2, % humidity, 37°C sterile incubator for 3-5 days.

On the day of experiment, cells were pretreated with 80uM WIN or its vehicle (0.5% DMSO in sterile PBS) for 30 minutes and then treated with 4uM forskolin for 1 hour. Following cell lysis, a cAMP assay (ABCAM, Cat# ab65355) was performed per manufacturer specifications. As no sex differences were observed, data from both sexes have been pooled.

### 4.8 Gut Motility Assay

Twenty minutes after 3mg/kg IP WIN, animals were given 0.25mL black marker (10% charcoal in 5% gum Arabic) via gavage. 20-30 minutes post-gavage, animals were sacrificed via decapitation and fresh GI tract and small intestine were removed. The distance black marker traveled from pylorus was recorded and expressed as a ratio between distance traveled and total length of small intestine from pylorus to caecum [75].

### 4.8 Eye Wipe Test

Capsaicin (0.1% in 0.01% EtOH) (Sigma-Aldrich, Cat#M2028) was placed in the corner of the animal’s eye. Time spent wiping over 2 minutes was recorded [76].

### 4.9 Surgical Procedures

Spared nerve injury was used as a model of neuropathic pain [77, 78]. In brief, mice were anesthetized under isoflurane anesthesia (5.0% induction, 2.5 % maintenance) and the skin and muscle of the left thigh were incised and the sciatic nerve along with its three branches (common peroneal, tibial and sural) was exposed. A tight ligature using a 5-0 silk suture was placed around the tibial and common peroneal branches, taking care to not stretch or damage the sural nerve, after which the nerve distal to the ligature was transected. The skin was closed using an auto clip and mice were returned to their home cages to recover. If an animal was insensate after surgery (ie unsuccessful surgery), it was excluded from experiment. Sham surgeries were done identically to the SNI surgery; however, no portion of the sciatic nerve was ligated or transected.

### 4.10 Behavior

All experiments used mice of both sexes and were performed during the animals’ light cycle, between 0900h and 1200h. White noise was playing in the behavior room during testing. Experimenters were blinded to genotype and surgical, inflammatory, or treatment state. The same mice were used for von Frey, Hargreaves, acetone test, paw volume, and paw temperature within the separate behavior paradigms.

### 4.11 Von Frey

In instances where mechanical hypersensitivity was tested, the up-down method of Chalpain et al was used [79]. In brief, animals were allowed to acclimate to plexiglass boxes with mesh flooring for at least one hour before testing. The plantar surface of the hindpaw was stimulated with a series of Von Frey filaments (Stoelting, Cat#58011). If an animal responded to the stimulus by lifting or shaking the paw, spreading the toes, or licking or grooming the paw, the next lowest filament was used to stimulate the hindpaw; if no response was seen, the next highest filament was tested. This continued until the animals were either tested five times or reached the highest filament. The mechanical withdrawal threshold was calculated as the weight at which the animals responded 50% of the time. In neuropathic pain experiments, only the lateral aspect of the hindpaw was tested.

### 4.12 Hargreaves

In instances where heat sensitivity was tested, the Hargreaves test was used [80]. In brief, animals were allowed to acclimate to the Hargreaves apparatus (IITC, part #336G). After acclimation, a laser was directed to a section of the apparatus corresponding to the plantar surface of the hindpaw. Intensity was set so that animals consistently baselined at 8-15 seconds. To avoid tissue damage, the laser was set to cut off at 25 seconds. Withdrawal latencies were defined as the amount of time it took an animal to remove its paw from the heated surface.

### 4.13 Intraplantar Inflammation

As a measure of inflammation after local carrageenan, paw volume was recorded using water displacement via plethysmometer (IITC, part #520MR) [54]. Additionally, forward-looking infrared camera (FLIR) (model T650SC, Wilsonville, OR) was used to measure paw temperature [54]. Image analysis of the plantar surface was performed using the FLIR ResearchIR Max 4 software.

### 4.14 Tail Flick

In brief, animals were restrained in a clean towel, and a laser (IITC, part #336G) was focused on the midpoint of the tail. The withdrawal latency was defined as the time it took for the animal to remove its tail from the heat source. Intensity was set so that animals consistently baselined at 5-8 seconds. To prevent tissue damage, the maximum amount of time animals were exposed to the heat source was 15 seconds.

### 4.15 Acetone Test

In instances where cold sensitivity was tested, the acetone test was used [78]. In brief, 100% acetone (Fisher, Cat#AI6P-4) was applied to the lateral aspect of the plantar hindpaw. Withdrawal latencies are defined as the amount of time in seconds the animal spent shaking, grooming, or raising its paw was recorded over the course of one minute.

### 4.16 Graphics

In figures that used experimental schematic graphics, BioRender.com was used to generate panels. All graphics were exported under a paid license.

### 4.17 Statistical Analysis

Prism 10 software (GraphPad, San Diego, CA, USA) was utilized to generate all graphs and statistical analysis. Single comparisons were performed using Student’s t-test, and multiple comparisons were performed using a one-way or two-way ANOVA with Bonferroni post hoc tests for cross-group comparisons. Relevant statistics are included in figure legends. All data are represented as the standard error of the mean (SEM). A p-value of <0.05 was used to determine statistical significance. All statistical values and corresponding tests can be found in Tables 2-24.

## Acknowledgements

The authors would like to thank Aspen Samuel for technical expertise and current and past members of the Neuroimmunology and Behavior laboratory for their contributions.

## Author Contributions

MDB and ALM conceptualized the study. MDB, CMC, and JKE provided resources and materials for the study. ALM, ARG,KMG, TAS-P, LRB, and DMJ contributed to the experimental design and data acquisition, analysis, and interpretation. ALM, ARG, KMG, and drafted and edited the manuscript and figures.

---

**Supplemental Fig. 1:** Sex and dose do not influence the inability of CB1R restricted to peripheral nociceptors to mediate reflexive analgesia. (A) Animals responded to WIN independent of sex or dose Dunnett’s multiple comparisons test; n=5 animals, \*\*\*\**p*<0.0001 vs. baseline). (B) A low dose (1mg/kg) of WIN induced analgesia in WT, but not CB1R^LOXTB^ or Na_v_1.8^+/-^:CB1R^LOXTB^ animals (Tukey’s multiple comparisons test, n=5-8 animals, \*\**p*<0.01 vs. baseline). (C) A high dose (3mg/kg) of WIN induced analgesia in WT, but not CB1R^LOXTB^ or Na_v_1.8^+/-^:CB1R^LOXTB^ animals (Tukey’s multiple comparisons test; n=5-7 animals, \*\*\**p*<0.001). Dotted lines on graphs indicate baseline values (5-8 seconds). Grey boxes indicate 10-, 40-, and 80-minute (A) or 15- and 60-minute (B and C) time points.

**Supplemental Fig. 2:** Generation of Na_v_1.8^+/-^:CB1R^fl/fl^ animals. (A) RNAscope *In Situ* hybridization of *Cnr1* (magenta) and *Scn10a* (green) mRNA in the lumbar (L4-6) DRGs. Scale bar = 20 μm. (B) Average colocalized *Cnr1* and *Scn10a* puncta (Tukey’s multiple comparisons test; n=5-6 animals, \**p*<0.05 WT vs. Na_v_1.8^+/-^:CB1R^fl/fl^). Cell size distribution of *Cnr1*, *Scn10a*, and colocalized *Scn10a* puncta positive cells in WT (C) and Na_v_1.8^+/-^:CB1R^fl/fl^ (D) lumbar DRGs.

**Supplemental Fig. 3:** Area Above the Curve Effect Size for Na_v_1.8^+/-^:CB1R^fl/fl^ Inflammatory Pain Model. (A) WIN caused analgesia in response to mechanical stimuli in both CB1R^fl/fl^ and Na_v_1.8^+/-^:CB1R^fl/fl^ animals (Sidak’s multiple comparison’s test, n=8-12 animals, \*\*\*\**p*<0.0001, \*\**p*<0.01, \**p*<0.05). (B) WIN caused analgesia in response to thermal stimuli in both CB1R^fl/fl^ and Na_v_1.8^+/-^:CB1R^fl/fl^ animals (Sidak’s multiple comparison’s test, n=8-12 animals, \*\**p*<0.01, \**p*<0.05).

