## Supplementary figures and images for "Nociceptor-restricted cannabinoid receptor 1 contributes to chronic but not acute analgesia"

### Supplemental Figure 1

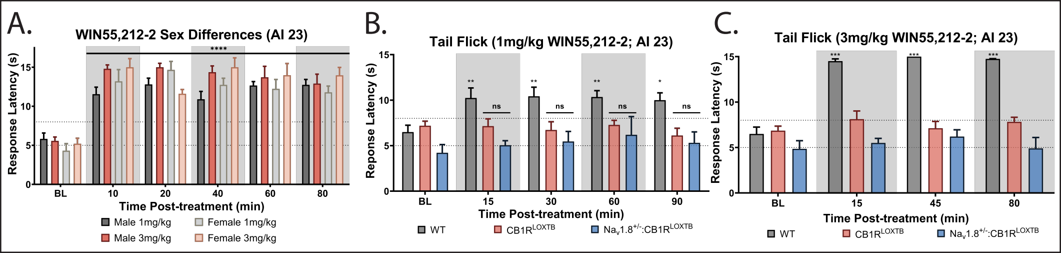

### Supplemental Figure 2

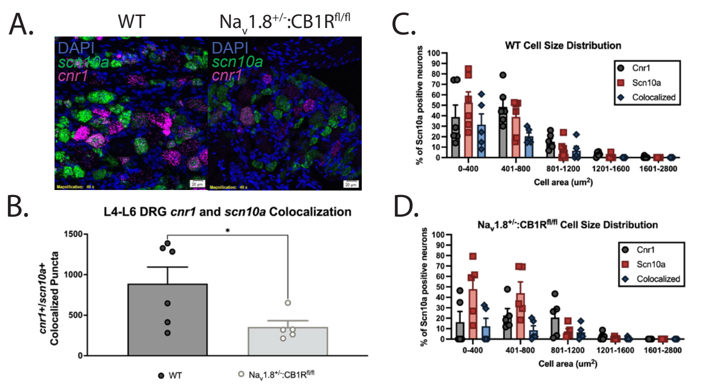

### Supplemental Figure 3

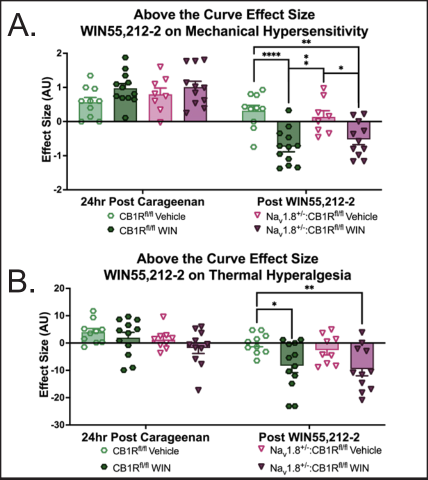
